# *echolocatoR*: an automated end-to-end statistical and functional genomic fine-mapping pipeline

**DOI:** 10.1101/2020.10.22.351221

**Authors:** Brian M. Schilder, Jack Humphrey, Towfique Raj

**Affiliations:** Nash Family Department of Neuroscience & Friedman Brain Institute, Icahn School of Medicine at Mount Sinai, New York, NY, United States of America; Ronald M. Loeb Center for Alzheimer’s disease, Icahn School of Medicine at Mount Sinai, New York, NY, United States of America; Department of Genetics and Genomic Sciences, Icahn School of Medicine at Mount Sinai, New York, NY, United States of America; Icahn Institute for Data Science and Genomic Technology, Icahn School of Medicine at Mount Sinai, New York, NY, United States of America; Estelle and Daniel Maggin Department of Neurology, Icahn School of Medicine at Mount Sinai, New York, NY, United States of America

**Keywords:** Genome-wide association, Expression quantitative trait loci (eQTL), Neurogenetics, Bioinformatics, Fine-mapping

## Abstract

**Summary:** *echolocatoR* integrates a diverse suite of statistical and functional fine-mapping tools in order to identify, test enrichment in, and visualize high-confidence causal consensus variants in any phenotype. It requires minimal input from users (a summary statistics file), can be run in a single R function, and provides extensive access to relevant datasets (e.g. reference linkage disequilibrium (LD) panels, quantitative trait loci (QTL) datasets, genome-wide annotations, cell type-specific epigenomics, thereby enabling rapid, robust and scalable end-to-end fine-mapping investigations.

**Availability and implementation:** *echolocatoR* is an open-source R package available through GitHub under the MIT license: https://github.com/RajLabMSSM/echolocatoR

## 1. Introduction

Genome-wide association studies (GWAS) across a variety of phenotypes and quantitative trait loci (QTL) have identified many significant genetic associations. However, widespread non-independence between genomic variants due to linkage disequilibrium (LD) makes it difficult to distinguish causal variants from correlated non-causal variants (Pasaniuc and Price, 2017; Pritchard and Przeworski, 2001; Yang *et al.*, 2011). Fine-mapping aims to identify the causal variant(s) and thus the mechanisms underlying a phenotype (Spain and Barrett, 2015; Trynka *et al.*, 2015). This methodology has been especially important to the study of medical conditions such as diabetes (Gaulton *et al.*, 2015; Mahajan *et al.*, 2018), rheumatoid arthritis (Kichaev and Pasaniuc, 2015; Westra *et al.*, 2018), and obesity (Zhang *et al.*, 2018).

Many fine-mapping tools have been developed over the years (Spain and Barrett, 2015; Trynka *et al.*, 2015), each of which can nominate partially overlapping sets of putative causal variants. It can therefore be useful to compare results from multiple fine-mapping methods with complementary strengths and weaknesses, such as the ability to model multiple causal variants or incorporate functional annotations. However, these powerful methods are underutilized in no small part due to technical reasons (e.g. not available in the same programming language, idiosyncratic file inputs/outputs, gathering and formatting of datasets). We therefore developed *echolocatoR*, an open-source R package that conducts end-to-end statistical and functional fine-mapping, annotation, enrichment and plotting that only requires GWAS/QTL summary statistics as input (Fig. 1a).

**Figure 1.**
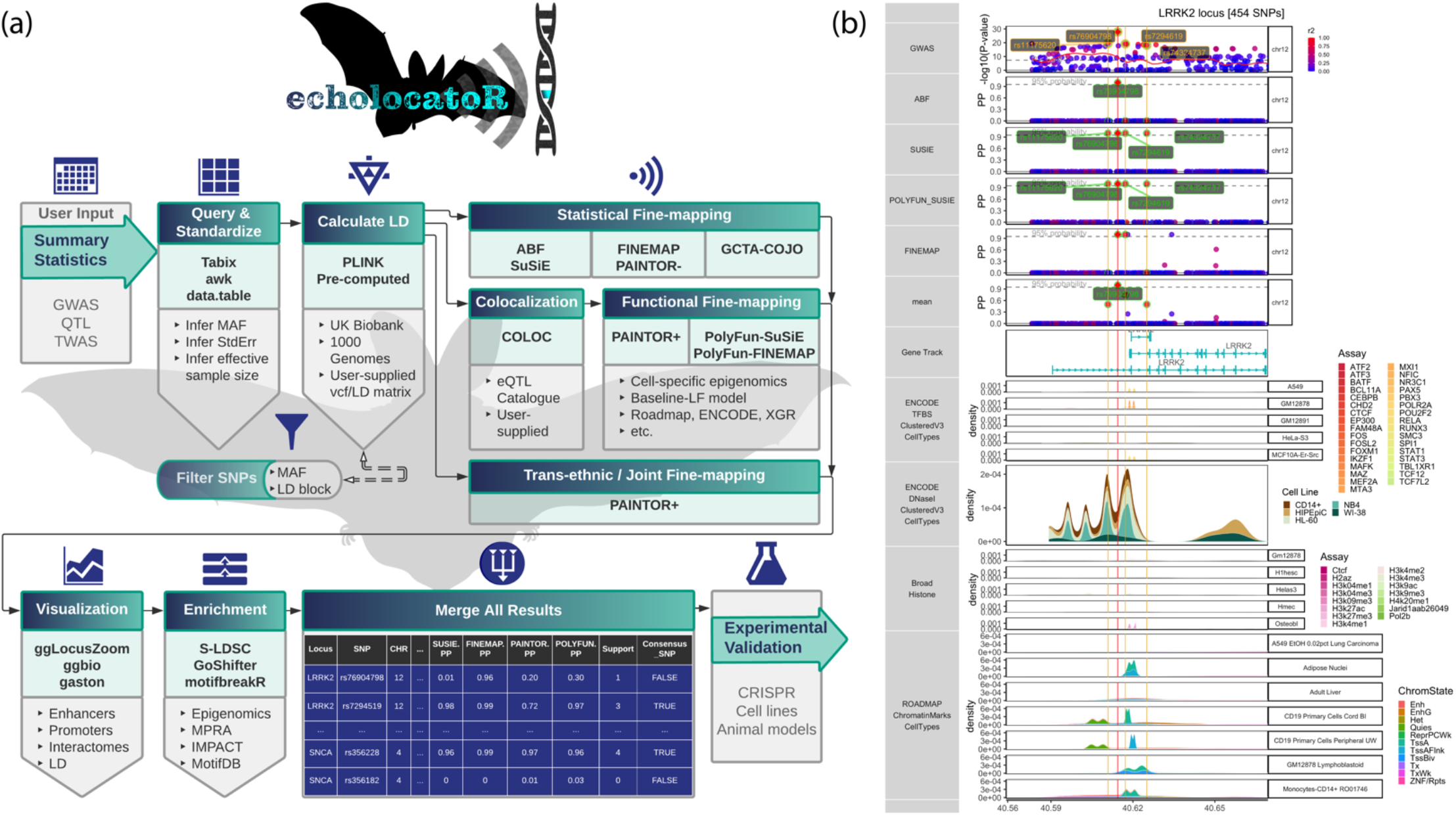
*echolocatoR* facilitates automated end-to-end fine-mapping. (**a**) Workflow of the *echolocatoR* pipeline: 1) user specifies the path to their full GWAS/QTL summary statistics, 2) locus subsets are queried and saved in a standardized format, 3) LD is extracted, computed from VCF, or supplied by the user, 3) statistical, functional, and/or trans-ethnic/joint fine-mapping are performed, 4) locus-specific fine-mapping results and selected annotations are visualized in tracks, 5) enrichment tests can be performed on different SNP groups using the various available annotations (see Section 2.2), 6) GWAS/QTL data, fine-mapping results and annotations are merged into a file with one SNP-row, 7) narrowed SNPs lists can be targeted in validation experiments. (**b**) Example multi-track plot for the PD locus LRRK2: 1) Manhattan plot of GWAS p-values (gold labels and vertical lines indicate Consensus SNPs), 2- 5) tool-specific fine-mapped posterior probability (PP) with 95% credible set (CS_95%_) SNPs labeled (green labels), 6) mean per-SNP PP across all fine-mapping tools, 7) gene transcript models, 8) transcription factor binding site (TFBS) annotations from ENCODE, 9) cell-type-specific histone modifications from ENCODE (Broad Institute, Bernstein lab), 10) cell-type-specific chromatin marks from Roadmap. Vertical red lines indicate the location of the lead GWAS SNP.

## 2. Implementation

The full *echolocatoR* fine-mapping pipeline can be run using just the *finemap_loci()* function, which ultimately produces an organized folder structure containing study- and locus-specific multi-tool fine-mapping results tables and annotated multi-track plots. If some stage of the pipeline has been run previously for a given locus, *finemap_loci()* will automatically detect and use the associated files, saving time for when testing different parameters. Most *echolocatoR* functions can run on a standard laptop (tested on a MacBook Pro with a 2.3 GHz Intel Core i5 processor and 8 GB 2133 MHz LPDDR3 memory), or take full advantage of its parallelizing capabilities on a high performance computing (HPC) cluster.

### 2.1. Rapid, robust, and scalable fine-mapping

By default, *echolocatoR* automatically indexes the user’s summary statistics file using Tabix (Li, 2011) for rapid on the fly querying. Locus-specific summary statistics are then extracted, standardized, and filtered according to user-controllable parameters such as window size (± 1Mb surrounding the index SNP by default), minor allele frequency (MAF) threshold, LD block, and many other features.

*echolocatoR* integrates a suite of existing fine-mapping tools, which currently includes: ABF (Benner *et al.*, 2016; Wellcome Trust Case Control Consortium *et al.*, 2012; Wakefield, 2007), GCTA-COJO (Yang *et al.*, 2012), FINEMAP (Benner *et al.*, 2016), SuSiE (Wang *et al.*, 2018), PolyFun (Weissbrod *et al.*, 2019), and PAINTOR (Kichaev *et al.*, 2017), the latter of which can be run with (i.e. PAINTOR+) or without (PAINTOR-) functional annotations. Colocalization tests between pairs of GWAS and/or QTL can also be performed using *coloc* (Giambartolomei *et al.*, 2014) to identify locus-specific phenotype-relevant tissues and cell types and prioritize GWAS/QTL datasets for joint functional fine-mapping.

Each fine-mapping tool produces its own 95% Credible Set (CS_95%_). The precise meaning of this term varies by tool but can be understood as the SNPs with 95% probability of being causal in the phenotype of interest. However, inter-tool comparisons have observed that there is substantial heterogeneity in their CS_95%_ (see (Weissbrod *et al.*, 2019)), leading to questions about the validity of any single tool in all situations, which can be strongly influenced by the degree of LD complexity and the true number of causal SNPs (Pasaniuc and Price, 2017; Pritchard and Przeworski, 2001; Yang *et al.*, 2011). We therefore define Consensus SNPs as those that were identified in the CS_95%_ of two or more tools, representing high-confidence putative causal SNPs. Indeed, we have shown that these Consensus SNPs have significantly higher predicted regulatory impact than either index SNPs or individual tool CS_95%_ SNP sets in Parkinson’s Disease (PD) (Schilder and Raj, 2020). *echolocatoR* automatically adds columns for Support (the number of tools that a given SNP was in the CS_95%_), Consensus SNP status, as well as mean posterior probabilities (PP) across all fine-mapping tools used.

### 2.2. Extensive database access

#### 2.2.1. Linkage disequilibrium

A common barrier to performing accurate fine-mapping is access to the appropriate LD reference panels. Currently, application programming interface (API) access is provided for 1000 Genomes Phases 1 & 3 (with selectable subpopulations) (Consortium and The 1000 Genomes Project Consortium, 2015), UK Biobank (Bycroft *et al.*, 2018; Sudlow *et al.*, 2015; Weissbrod *et al.*), or user-supplied VCF files or LD matrices. Unlike existing LD querying tools (Machiela and Chanock, 2015), *echolocatoR* does not restrict the size of LD matrices to allow comprehensive fine-mapping of all loci regardless of size or complexity.

#### 2.2.2. Genome-wide annotations

Genome-wide annotations can be used to compute SNP-wise prior probabilities for functional fine-mapping (e.g. PolyFun, PAINTOR+). API access to a large compendium of genome-wide annotations and epigenomic data is provided, including: tissue and/or cell type/line-specific chromatin marks from Roadmap (Bernstein *et al.*, 2010; Satterlee *et al.*, 2019), ENCODE (Jou *et al.*, 2019), genic annotations through *biomaRt* (Durinck *et al.*, 2009),HaploReg (Zhbannikov *et al.*, 2017; Ward and Kellis, 2012), cell-type-specific epigenomic datasets(Nott *et al.*, 2019; Corces *et al.*), and hundreds of additional annotations through the R package *XGR* (http://xgr.r-forge.r-project.org/) (Fang *et al.*, 2016). *catalogueR*, another R package developed by our group, provides rapid API access to full summary statistics from 110 uniformly reprocessed QTL datasets (across 20 studies) with parallelized Tabix queries. *echolocatoR* can utilize all genome-wide annotations and datasets to compare enrichment across different SNP group (e.g. GWAS lead SNPs vs. CS_95%_ vs. Consensus SNPs) using XGR, GoShifter, and/or S-LDSC (Gazal *et al.*, 2017; Finucane *et al.*, 2015; Bulik-Sullivan *et al.*, 2015).

#### 2.2.3. *In-silico* validation

We also built in API access to *in silico* validation datasets, including massively parallel reporter assays (MPRA) (van Arensbergen *et al.*, 2019; Tewhey *et al.*, 2018), S-LDSC heritability enrichment, and predictions from multiple machine learning models trained on tissue- and cell-type-specific epigenomic annotations: Basenji (Kelley *et al.*, 2018), and DeepSEA (Zhou and Troyanskaya, 2015) (provided by (Dey *et al.*)) as well as IMPACT (Amariuta *et al.*, 2019). Lastly, we integrated *motifbreakR* which uses a comprehensive set of algorithms and position weight matrices (n = 9,933) to assess whether fine-mapped variants fall within sequence motifs and to what extent they disrupt binding to specific transcription factors (Coetzee *et al.*, 2015).

### 2.3 Multi-track plotting

High-resolution multi-track plots are automatically generated for each locus (Fig. 1b) and can include any combination of the following tracks: Manhattan plots of GWAS/QTL P-values or tool-specific fine-mapping posterior probabilities (PP) colored by LD with the lead SNP, mean PP, gene body models, and all aforementioned genome-wide annotations. Plots can be further customized as returned *ggplot* objects.

## 3. Conclusion

Overall, *echolocatoR* removes many of the primary barriers to perform a comprehensive fine-mapping investigation while improving the robustness of causal variant prediction through multi-tool consensus and *in silico* validation using a large compendium of (epi)genome-wide annotations. Thus, we hope that *echolocatoR* will make fine-mapping a standard practice, thereby uncovering human disease etiology and accelerating the development of novel therapeutics.

## Supplementary information

Installation instructions (with an optional *conda* environment to minimize dependency conflicts), vignettes, example data, documentation of all functions and annotation datasets, as well as source code can be found in the *echolocatoR* website: https://rajlabmssm.github.io/echolocatoR

## Acknowledgements

We would like to thank Elisa Navarro, Gloriia Novikova, Cecilia Lindgren and Teresa Ferreira for their valuable feedback and suggestions. We would also like to thank Omer Weissbrod, Chris Glass, Alexi Nott for their guidance with data and/or tool integration. This work was supported in part through the computational resources provided by Scientific Computing at the ISMMS.

